# Coordinated Turning Behaviour of Loitering Honeybees (Apis Mellifera)

**DOI:** 10.1101/332379

**Authors:** Mandiyam Y. Mahadeeswara, Mandyam V. Srinivasan

## Abstract

Turning during flight is a complex behaviour that requires coordination to ensure that the resulting centrifugal force is never large enough to disrupt the intended turning trajectory. The centrifugal force during a turn increases with the curvature (sharpness) of the turn, as well as the speed of flight. Consequently, sharp turns would require lower flight speeds, in order to limit the centrifugal force to a manageable level and prevent unwanted sideslips. We have video-filmed honeybees flying near a hive entrance when the entrance is temporarily blocked. A 3D reconstruction and analysis of the flight trajectories executed during this loitering behaviour reveals that sharper turns are indeed executed at lower speeds. During a turn, the flight speed is matched to the curvature, moment to moment, in such a way as to maintain the centrifugal force at an approximately constant, low level of about 30% of the body weight, irrespective of the speed or the curvature of the turn. This ensures that turns are well coordinated, with few or no sideslips - as is evident from analysis of other properties of the flight trajectories.

## INTRODUCTION

It is a common experience that driving too fast around a corner can cause a car to skid or roll over; a passenger standing in a bus can tip over if the bus makes a turn at high speed; or an aircraft attempting to make a very tight turn can experience a sideslip. The reason is that the act of turning while simultaneously moving forward creates a centrifugal force that is directed away from the centre of curvature of the turn. Newtonian mechanics dictates that the magnitude of the centrifugal force is proportional to (a) the curvature (the reciprocal of the radius of the turn), and (b) the square of the speed (Anton, 1999). Hence, the sharper the turn, and the higher the speed, the greater the danger of losing control. Clearly, therefore, it makes sense to reduce one’s speed before commencing a turn, and to ensure that sharper turns are executed at a slower speed, in order to limit the centrifugal force to a safe and manageable value. This behaviour is adopted not only by car drivers, motorcyclists, bicyclists and runners, but also by several terrestrial and flying animals. Qualitative evidence to support such behaviour has been documented in race horses (Tan and Wilson, 2011), quolls (Wynn et al., 2015), houseflies (Wagner, 1986), fruitflies (Mronz and Lehmann, 2008) and bats (Aldridge, 1987). However, a quantitative analysis of the relationship between flight speed and curvature, and the implications for the resulting centrifugal force that is experienced during turns, have not yet been explored in any animal.

Fruitflies (*Drosophila*) flying in a contained environment display segments of straight flight, interspersed with saccadic turns (Fry, 2003; Muijres et al., 2015). These turns are executed by performing a pitch and a roll of the body axis, which together induce a rapid change in the direction of flight. Visually evoked escape maneuvers of fruit flies also include sharp turns (Muijres et al., 2014), which are much faster than the stereotyped body saccades. While these turns enable rapid, aggressive changes of flight direction, they are inevitably associated with sideslips arising from the high centrifugal force. It is of interest to enquire whether flying insects are also capable of performing turns that are coordinated in such a way as to prevent sideslips - for example, during loitering flight. In our study, we induce honeybees to loiter in front of their hive by blocking the entrance to the hive, which causes returning foragers to cruise in the vicinity ofthe hive entrance while they await entry. The behaviour of the bees in this ‘bee cloud’ is filmed using stereo video cameras and reconstructed in 3D to analyse their turning characteristics. The results reveal that loitering bees perform turns that are fully coordinated, and free of sideslips.

## METHODS

A non-captive honeybee colony (*Apis Mellifera*) was maintained on a semi-outdoor terrace on the rooftop of a building on the campus of the University of Queensland (St. Lucia). The bees were allowed to forage freely from the surrounding vegetation, without any restrictions. The experiment was commenced by temporarily blocking the hive entrance with a wooden strip (Fig. 1). The returning foraging bees were thus temporarily denied entry into the hive but flew near the vicinity of the hive entrance, making multiple attempts to gain access. The resulting ‘bee cloud’ was filmed using two synchronized digital cameras (Redlake), configured to obtain stereo data. The cameras recorded video at 60 fps with 500 × 500 pixel resolution.

**Figure 1:**
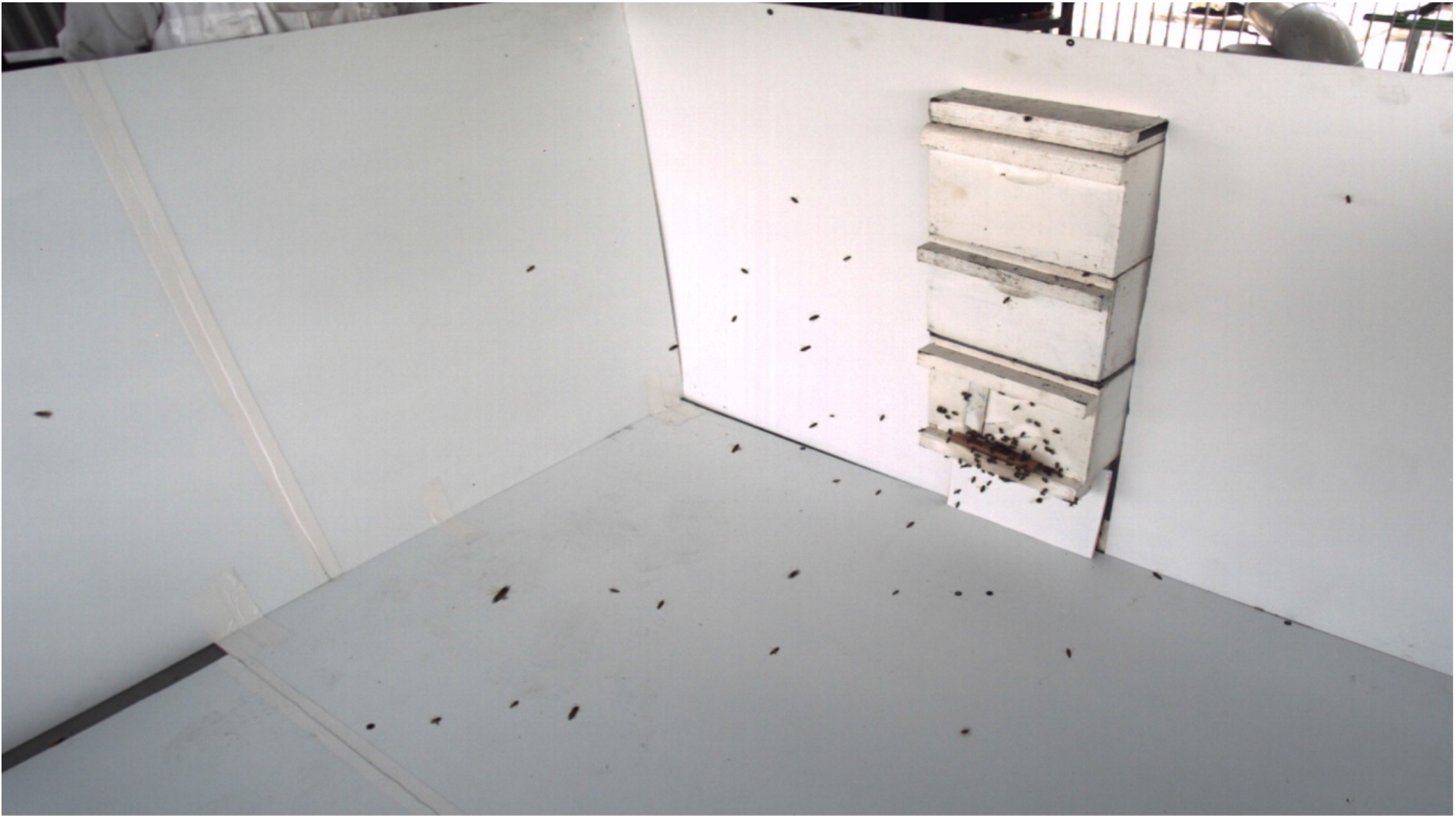
*A perspective view of the bee cloud*.

Before commencing the experiment, stereo camera calibration was performed to obtain the cameras’ intrinsic and extrinsic parameters. The video streams acquired by the two cameras were subsequently analysed by digitising the bee’s head and tail positions manually in each frame, to obtain the bee’s position coordinates in each view. A triangulation routine was executed to obtain the three-dimensional positional coordinates of each bee. The 3D coordinates of a bee were computable only when it was within the FOV of both cameras. The recording duration was 5.8 seconds (349 frames). The frames in the video footage carried varying numbers of bees, as individual bees entered or departed from the fields of view (FOV) of the two cameras.

The method used to compute the kinematic parameters of the flight were based on vector calculus (Anton, 1999). The tangential and normal components of the acceleration at any time instant ‘t’ can be computed as

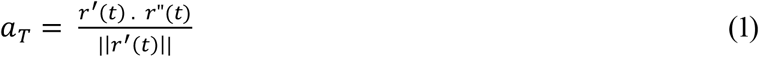

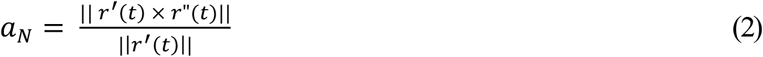

where *a_T_* ≜ tangential acceleration; *a_N_* ≜ normal or centripetal acceleration; *r*(*t*) ≜ position vector; *r*′(*t*) ≜ velocity vector and *r*″(*t*) ≜ total acceleration vector as function of time.

Mathematically, the curvature can be expressed as the rate of change of the unit tangent vector at a point. Using this concept, one can compute the magnitude of the curvature as function of time using the following vector algebra:

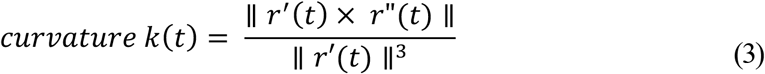

The radius of curvature (ρ) is expressed as the reciprocal of the curvature:

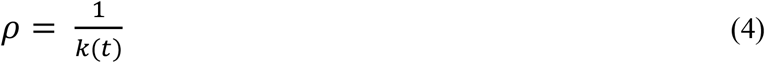

## RESULTS

The bee cloud data contains 3D position coordinates of a total of 66 bees. Fig. 1 shows a perspective view of the bee cloud at a particular instant of time. Fig. 2 shows the reconstructed 3D trajectories of the 66 bees, where each colour represents the trajectory of an individual bee. We used techniques of vector calculus to examine the flight characteristics of bees maneuvering in the cloud, by computing the following parameters:

a. The speed of each bee in the cloud, and its variation as a function of time
b. The acceleration of each bee in the cloud, and its tangential and centripetal components, and the variation of these parameters as a function of time
c. The curvature and radius of curvature of the flight trajectory, and its variation with time

**Figure 2:**
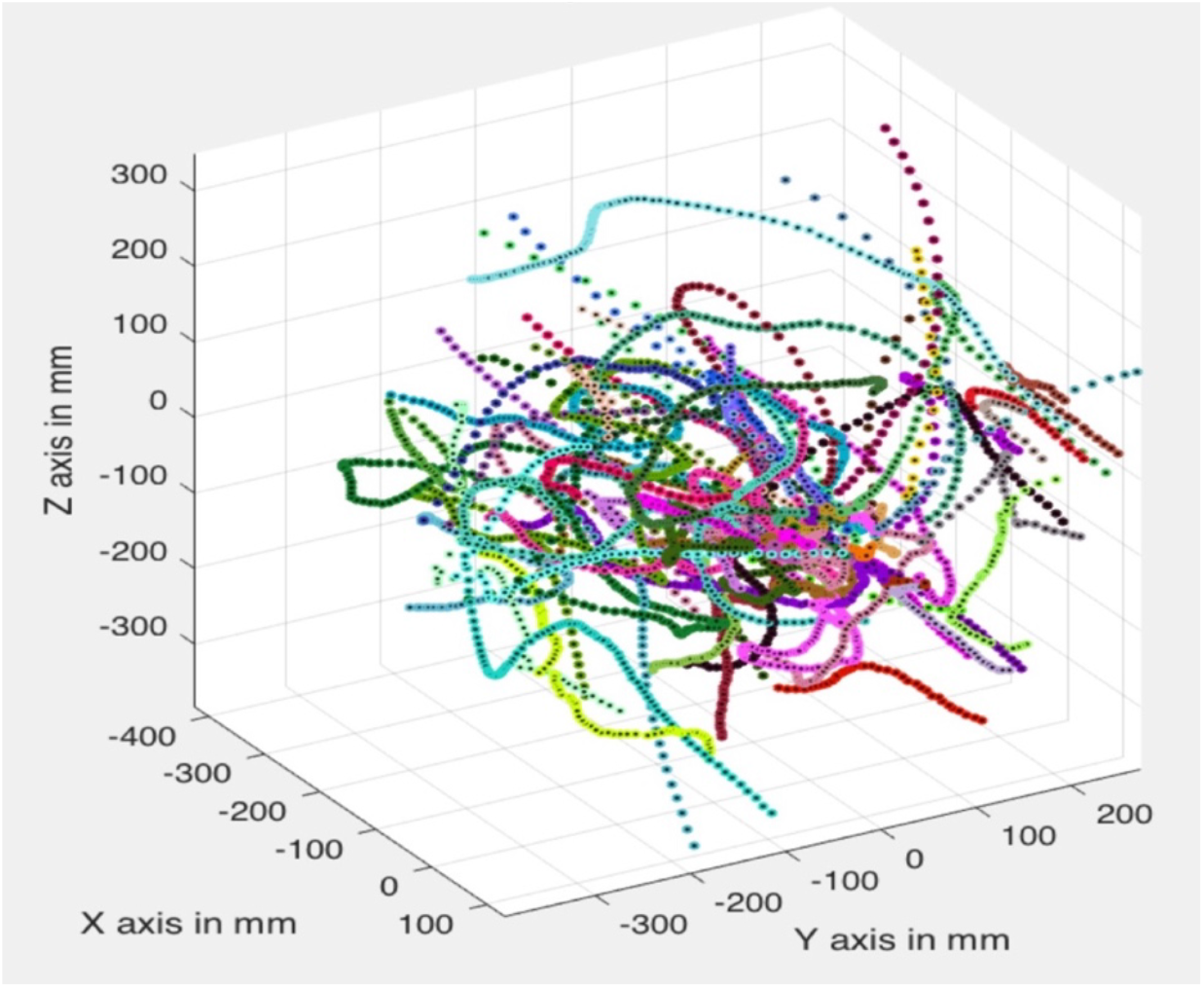
*Reconstructed 3D trajectories of 66 bees*.

The variation of each of the above parameters as a function of time is shown in Fig. 3 for a single bee.

**Figure 3(a-f):**
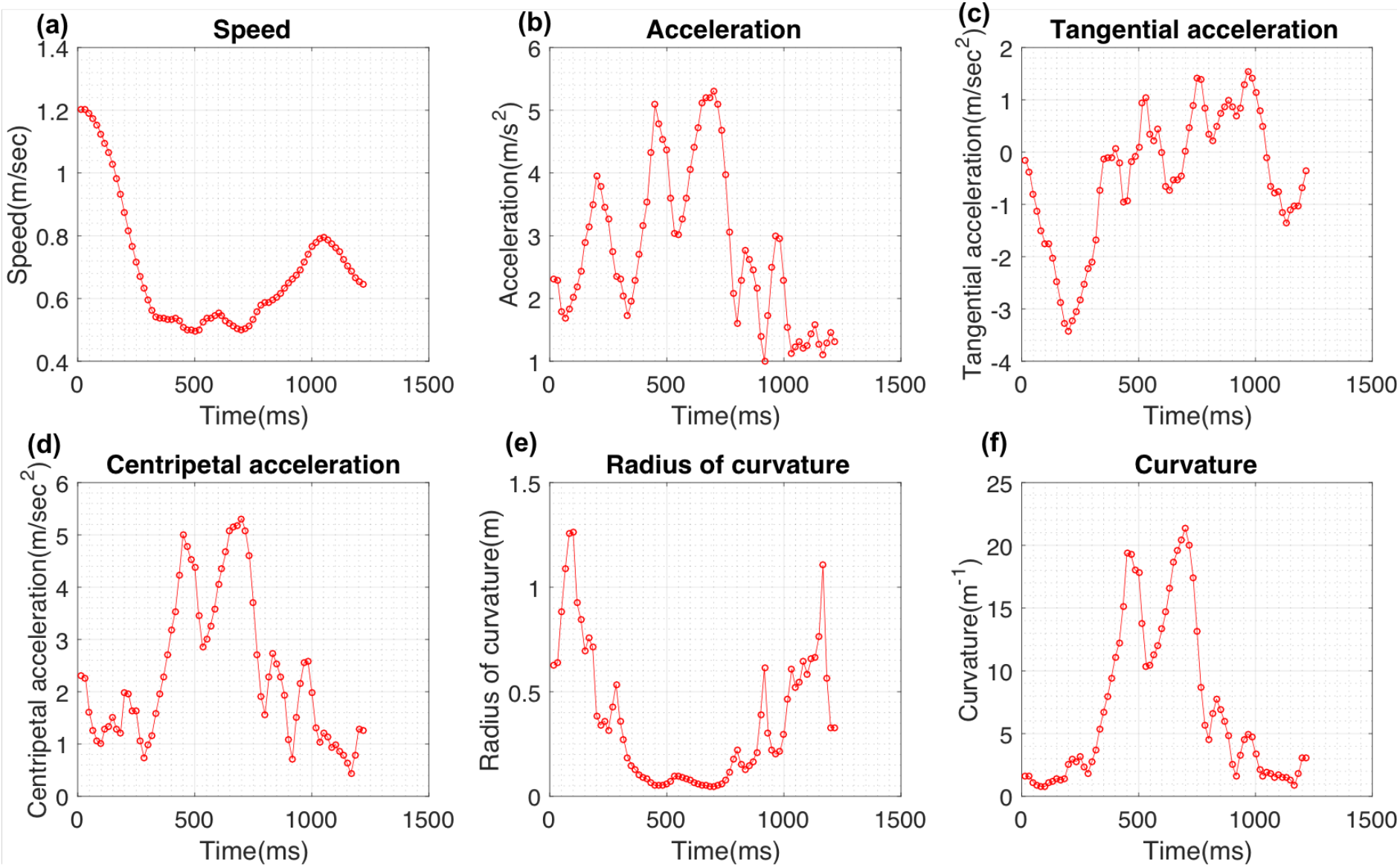
*Kinematic parameters of an individual bee in the cloud*

### General relationship between instantaneous speed, curvature and centripetal acceleration

In general, the speed of a bee varies continuously through its flight path, as shown in Fig. 3a for an individual bee. The mean speed of this particular bee is 0.45 m/sec. Certain bees exhibited high speeds, despite flying in close proximity to neighboring bees. For example, one individual reached a top speed of 2.61 m/s, while flying amidst 33 other bees in the cloud.

The average speed of all bees in the cloud was found to be 0.66 m/sec and the curvature of the trajectories displayed an average magnitude of 18 m^−1^. Average histograms of the speed and the curvature magnitudes of the trajectories of all 66 bees are shown in Figs. 4a and 4b, respectively. These histograms were obtained by computing an area-normalized histogram for each bee, and then averaging the results across the 66 bees.

**Figure 4(a):**
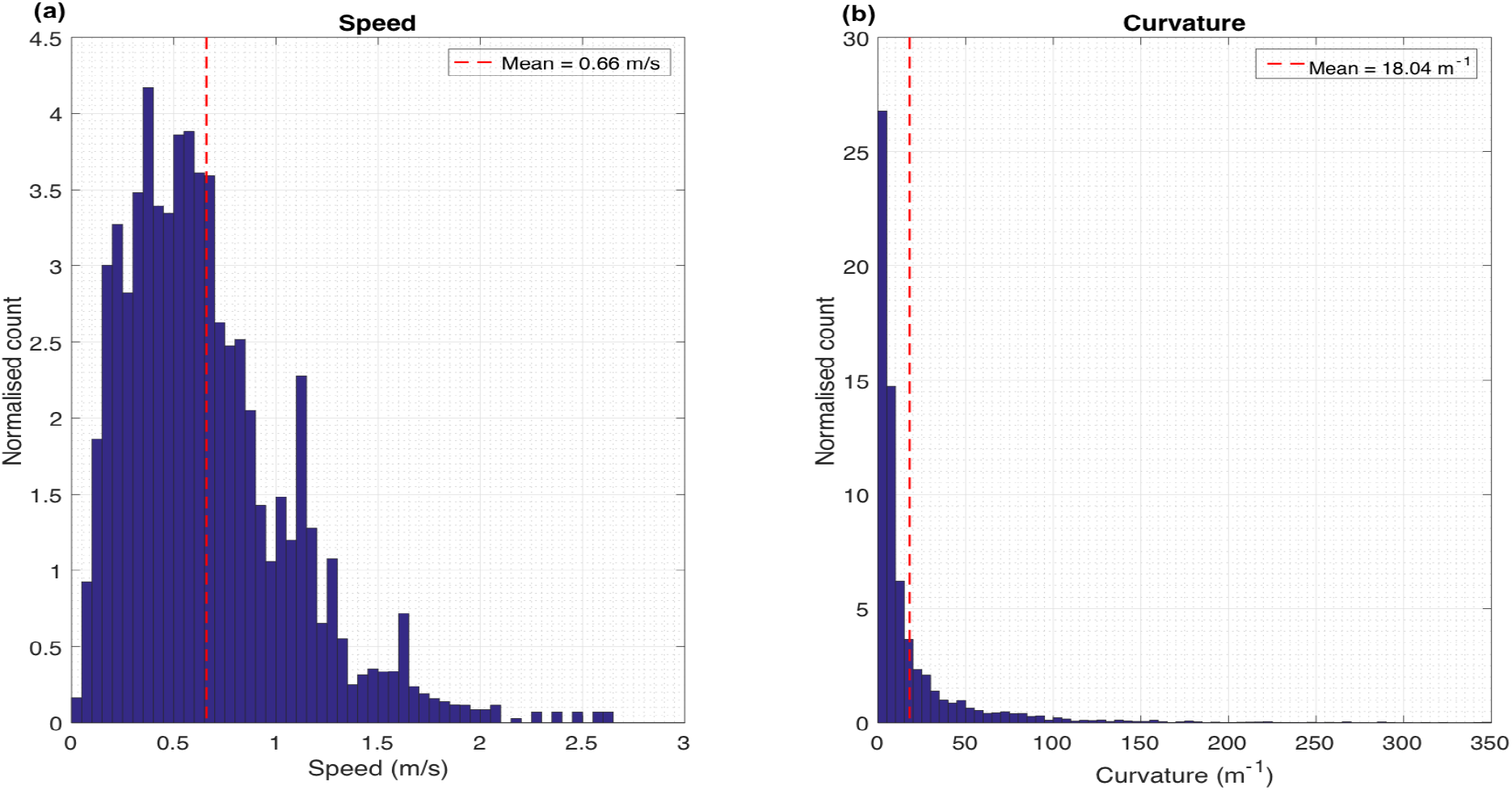
*Average histogram of speeds. (b): Average histogram of curvature magnitudes*.

The averaged histograms reveal a large variation in speed (ranging from 0 to 2.61 m/sec; Fig. 4a), as well as curvature magnitude (ranging from 0 to ~300 m^−1^; Fig. 4b). However, these histograms include the variations across all bees, which have different mean speeds and mean curvature magnitudes. A more representative measure of the average variability of speed and curvature within the trajectoiy of an individual bee is conveyed by the mean coefficient of variation (CV; averaged over all bees), which has a value of 0.32 for speed, and 1.5 for curvature magnitude.

Next, we calculated the tangential and centripetal components of the acceleration and plotted their variation as a function of time (Figs. 3c-d). The normalized histograms of tangential acceleration and the magnitude of the centripetal acceleration are shown in Figs. 5a-b respectively. The histogram of tangential acceleration clearly reveals that the flight contains acceleration and deceleration components, distributed approximately symmetrically about a value of zero (which corresponds to a constant tangential speed). The mean tangential acceleration, averaged across all bees, is 0.42 m/s^2^, which is not significantly different from zero (*p*=0.07; two tailed t-test). The mean standard deviation of the tangential acceleration is 2.0 m/s^2^. The magnitude of the centripetal acceleration, averaged across all bees, has a mean value of 2.80 m/s^2^, which is significantly different from zero (*p*=2.6×10^−25^; two tailed t-test). The mean CV of the centripetal acceleration magnitude is 0.51.

**Figure 5(a):**
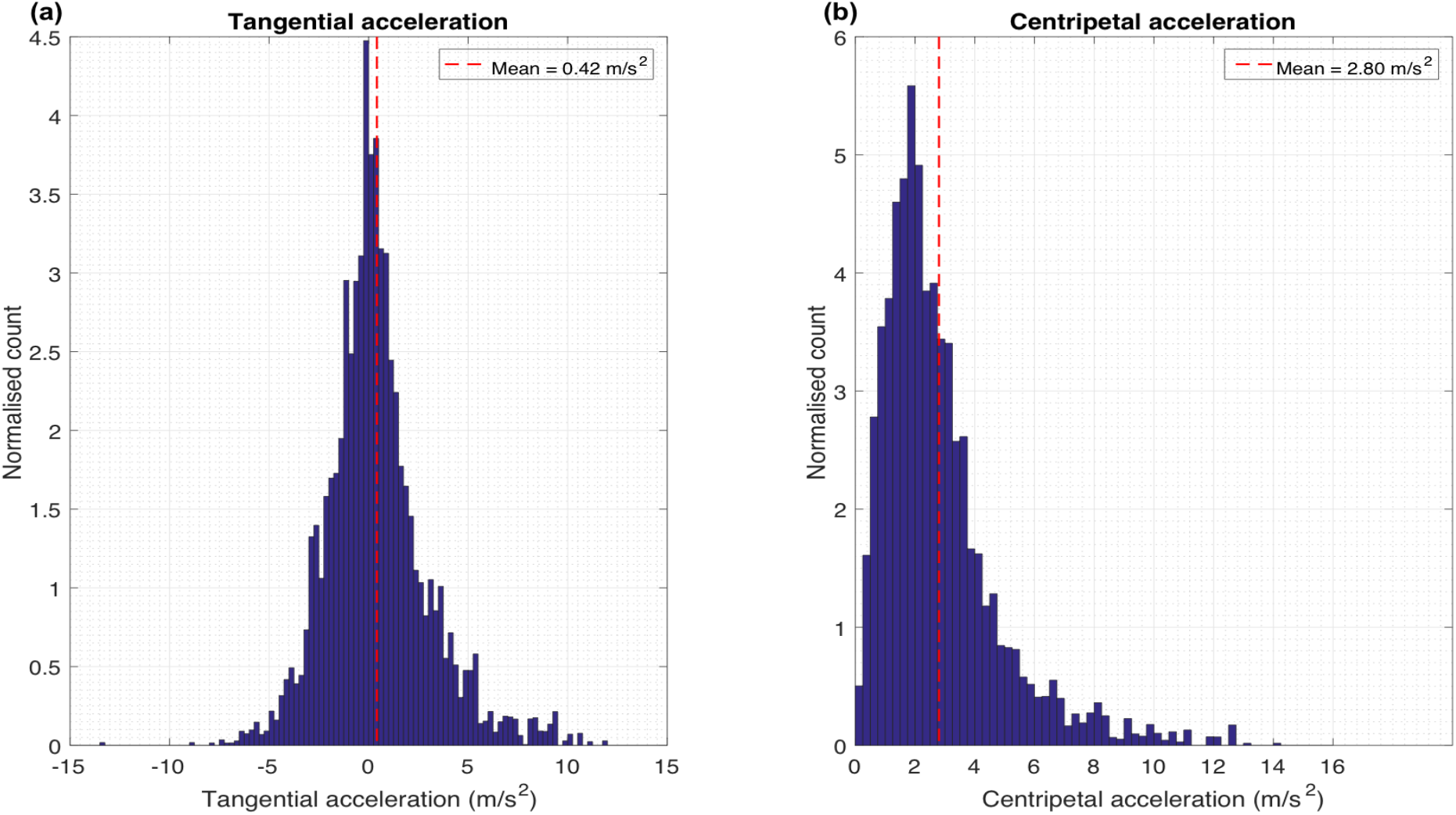
*Average histogram of tangential acceleration, (b): Average histogram of centripetal acceleration magnitudes*.

### Analysis of turning flights

The vector representations of the acceleration, and of its tangential and centripetal components, are shown in Fig. 6 for a segment of one trajectory. Inspection of these vectors reveals that the tangential acceleration is directed against the flight direction while entering the turn, and toward the flight direction while leaving the turn, indicating that the bee is decelerating while entering the turn and accelerating while leaving it (Fig. 6). On the other hand, the magnitude of the centripetal acceleration remains more or less constant through the turn. Thus, whereas the tangential component of acceleration varies dramatically during the flight (even changing its polarity), the magnitude of centripetal component of acceleration is more or less constant. This implies that the bee slows down while entering the turn, reaching a minimum speed close to the point of maximum curvature, and subsequently speeds up. The magnitude of the curvature, on the other hand, is low at the beginning of the turn, reaches a maximum value at the middle of the turn, and then declines toward zero as the turn is completed.

**Figure 6:**
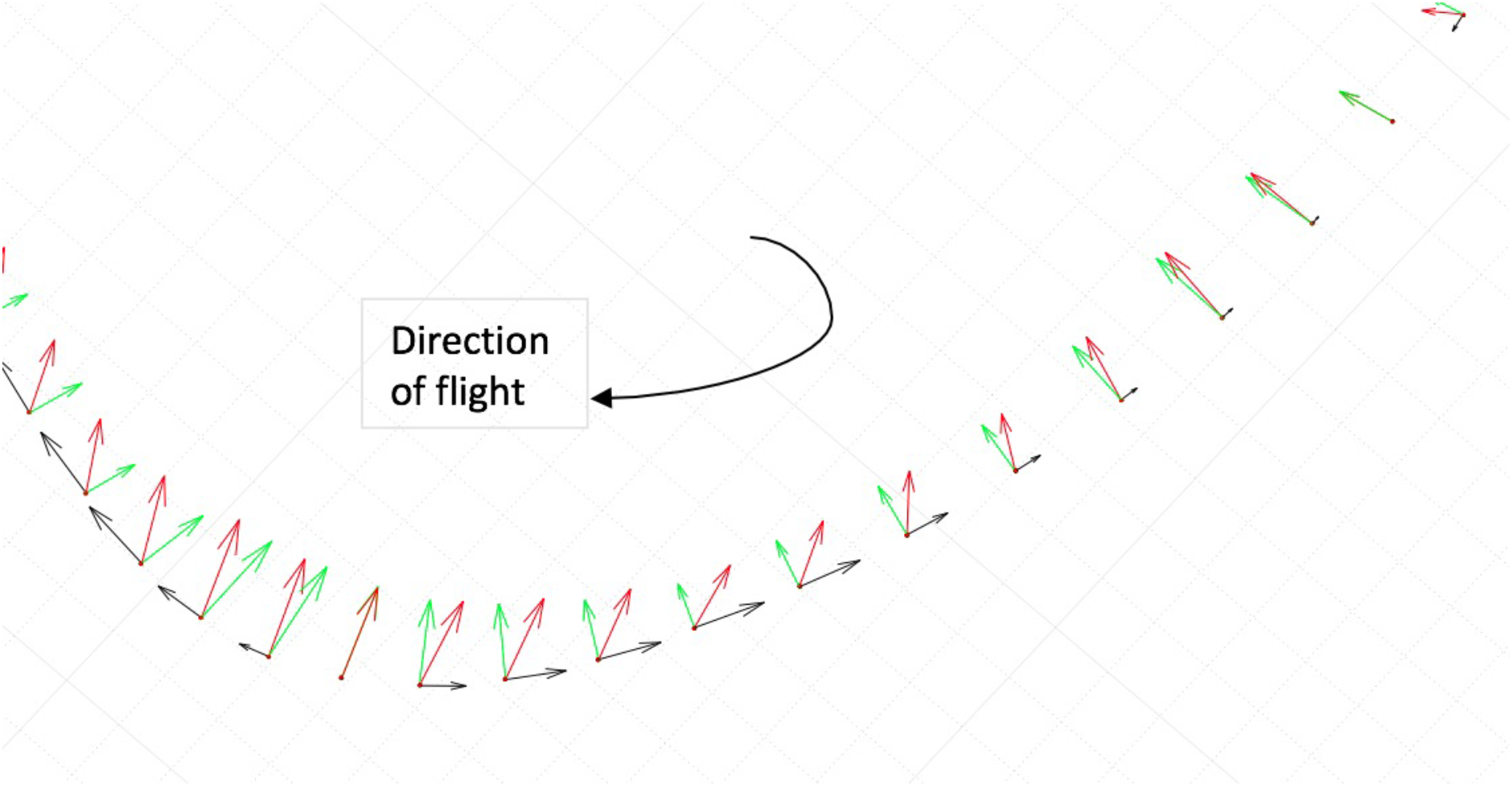
*Plan view of the acceleration and its components during a turn. The red, black, and green arrows represent the total, tangential and centripetal acceleration vectors respectively*.

This reciprocal variation of speed and curvature along the trajectory raises the possibility that the bees are maintaining their centripetal acceleration constant during the course of a turn. We examined the prediction of this hypothesis as follows.

We may write the centripetal acceleration as:

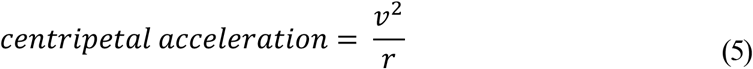

where ‘*r*’ is the instantaneous radius of curvature of the trajectory, and ‘*v*’ is the instantaneous bee speed. If the centripetal acceleration is constant, we have

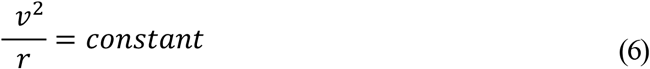

Therefore,

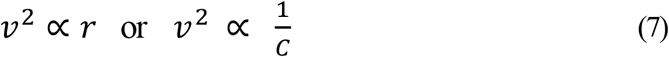

where *c* is the curvature of the trajectory, which is a measure of the sharpness of the turn.

If the bees are holding their centripetal acceleration constant (as hypothesised), then the following two predictions must hold:

a. a linear relationship between the radius of curvature and speed^2^.
b. an inverse relationship between curvature and speed^2^.

To test the hypothesis, we examined the variation of speed^2^ with the radius of curvature (ROC), and with the curvature of the trajectory for individual bees. These relationships are shown as scatterplots in Fig. 7. This data is plotted for a ROC range of 0.004 - 0.33 m, which corresponds to a curvature magnitude range of 3 – 250 m^−1^. Points along the trajectory at which the curvature magnitude was greater than 250 were not included in the plots, because the curvature measurements could be dominated by the errors in the image digitization process. Points at which the curvature magnitude was lower than 3, were considered to represent flight in an approximately straight line. These curvature limits were used to select the turning parts of the flight trajectory and exclude segments that corresponded to straight flight or very sharp turns. As a result of this process, the trajectories of two bees were removed from the total of 66 bee trajectories. Unless explicitly stated, the number of bees included in the subsequent analyses is 64.

**Figure 7(a-f):**
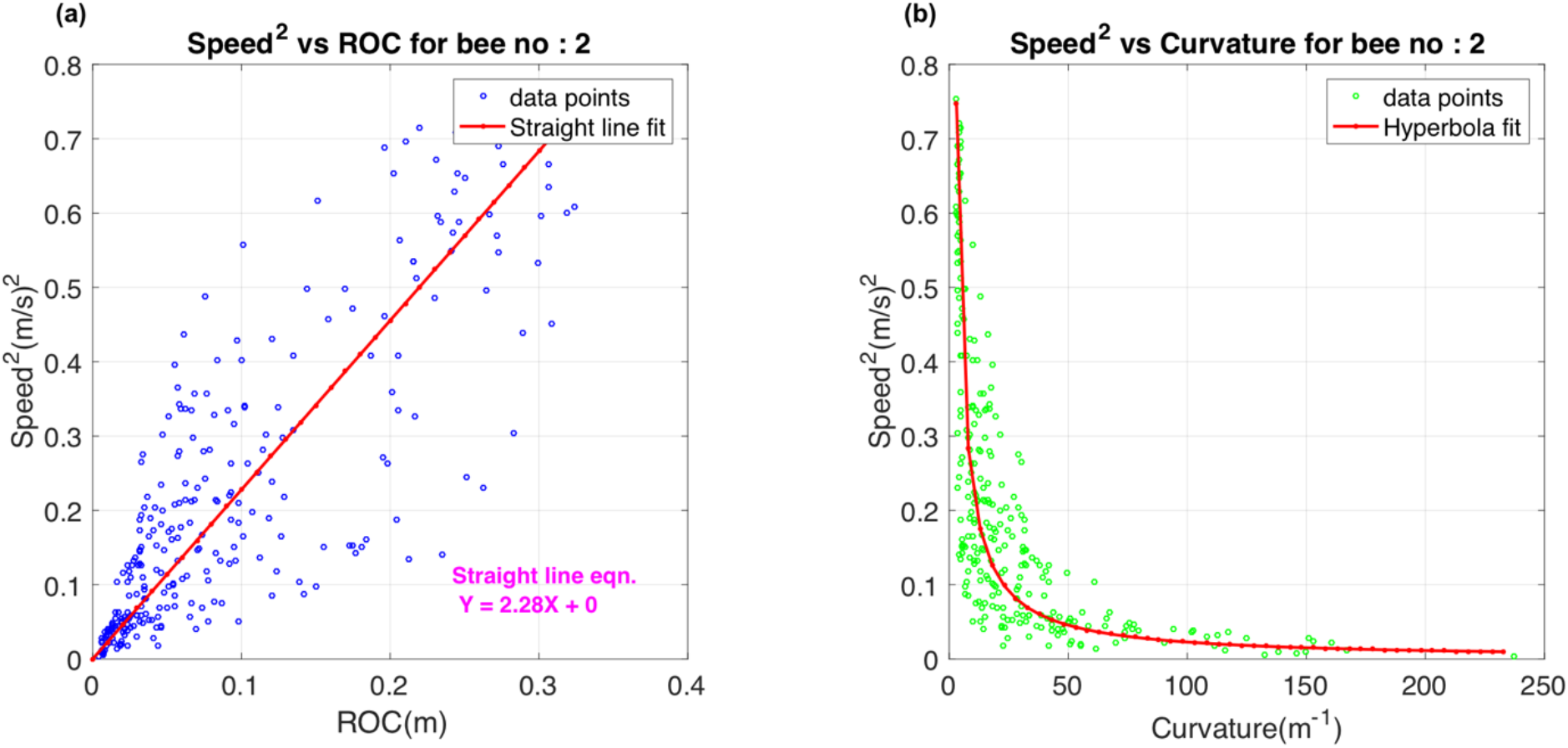
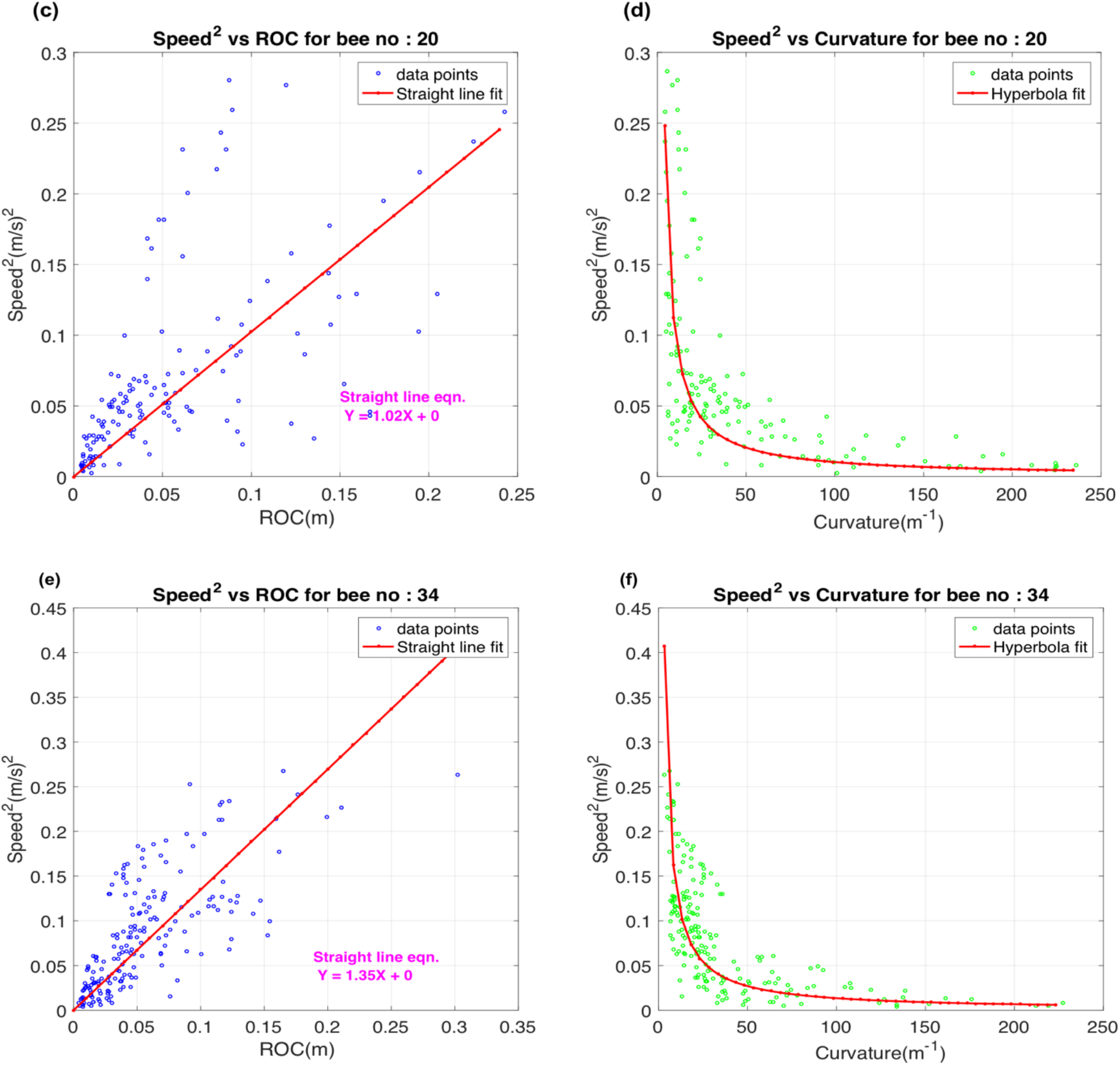
*Variation of speed^2^ with radius of curvature (ROC) and curvature for three individual bees*.

For the three scatterplots of speed^2^ versus ROC (Figs. 7a, c, e), we performed regression analysis on the data by forcing the ‘Y’ intercept to be zero and estimating the slope of the regression line. We used the ‘robust regression’ routine in Matlab, which removes outliers in the data. The estimated slope values were then used to generate theoretically predicted hyperbolic curves for the relationship between speed^2^ and curvature. Comparisons of the actual data with the theoretically predicted hyperbolic relationships are shown in Figs. 7b, 7d and 7f. The theory fits the data quite well for these three bees, corroborating the inverse relationship between curvature and speed^2^, as per our prediction. Additional examples for other bees are given in Section I of the Supplementary Material (SM).

The scatterplot of Fig. 8a illustrates the relationship between speed^2^ and ROC for all the bees. Each colour in the scatterplot represents a different bee. The scatterplot reveals that speed^2^ is approximately proportional to the radius of curvature, although the slope of the relationship varies from bee to bee. The overall slope of a linear regression, performed on the data in Fig. 8a, is 1.99. This implies that the magnitude of the centripetal acceleration is held at an approximately constant value of 1.99 m/s^2^. The scatterplot of Fig. 8(b) illustrates the relationship between speed^2^ and ROC for all the bees. The theoretically predicted hyperbolic relationship, obtained from the slope of the linear regression on the data of Fig. 8a, fits the data of Fig. 8b quite well.

**Figure 8(a):**
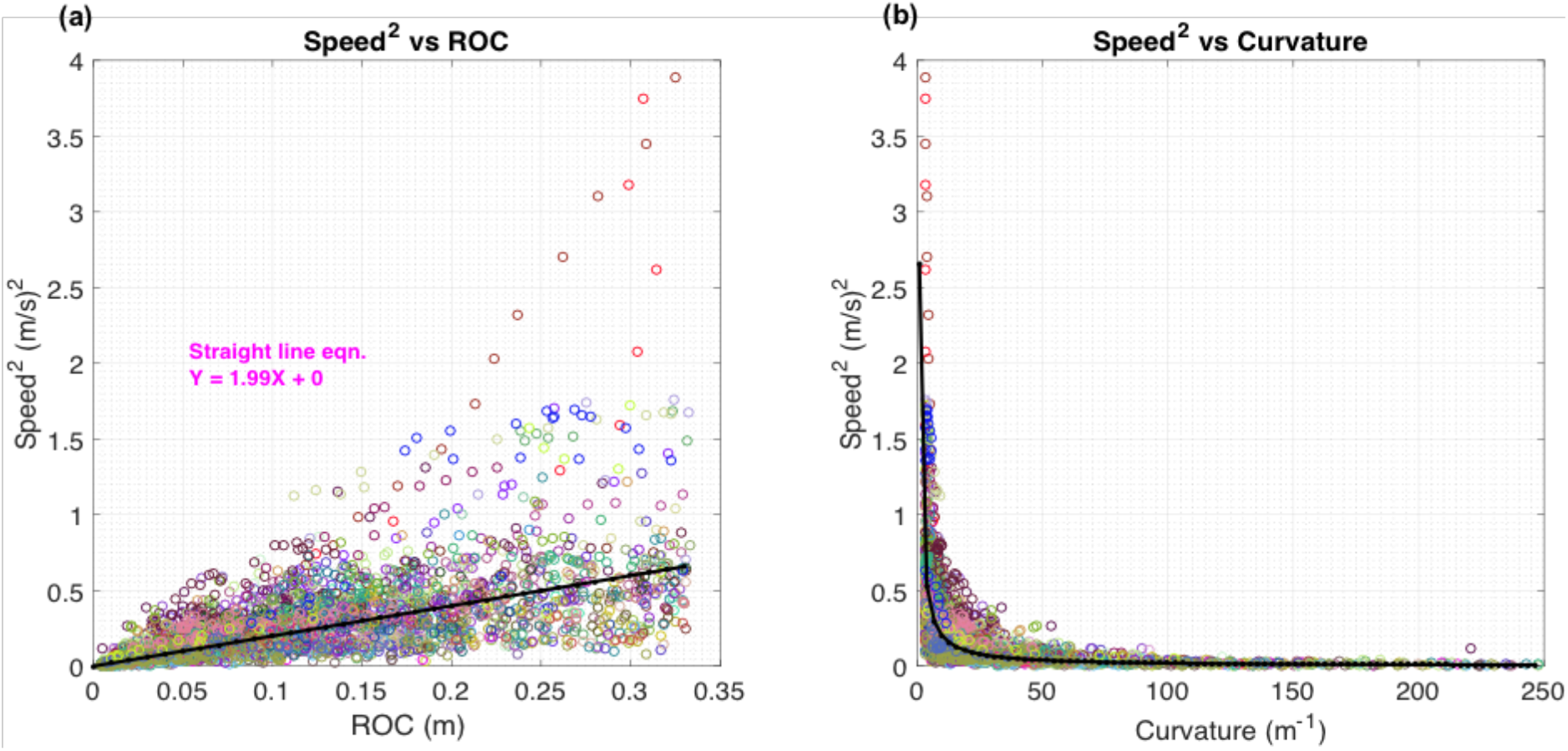
*Variation of speed^2^ with radius of curvature(ROC) (b): Variation of speed^2^ with curvature for 64 bee trajectories*.

To further test our hypothesis, we plotted the log-log relationship of speed^2^ versus radius of curvature (ROC) and speed^2^ versus curvature. As per our hypothesis, if there exists a linear relationship between speed^2^ and radius of curvature, then the relationship between log(speed^2^) and log(ROC) should be linear, with a slope of 1.0. If there exists an inverse relationship between speed^2^ and curvature, then the relationship between log(speed^2^) and log(curvature) should be again be linear, but with a slope of −1.

Fig. 9a shows the relationship between log(speed^2^) and log(ROC), plotted as a scattergram for the data pooled from the 64 bees. This relationship is approximately linear, with a slope of 0.9. This value is close to the value of 1.0 predicted by the hypothesis. The value of the regression slope for each individual bee is given in Section II of the SM.

**Figure 9(a):**
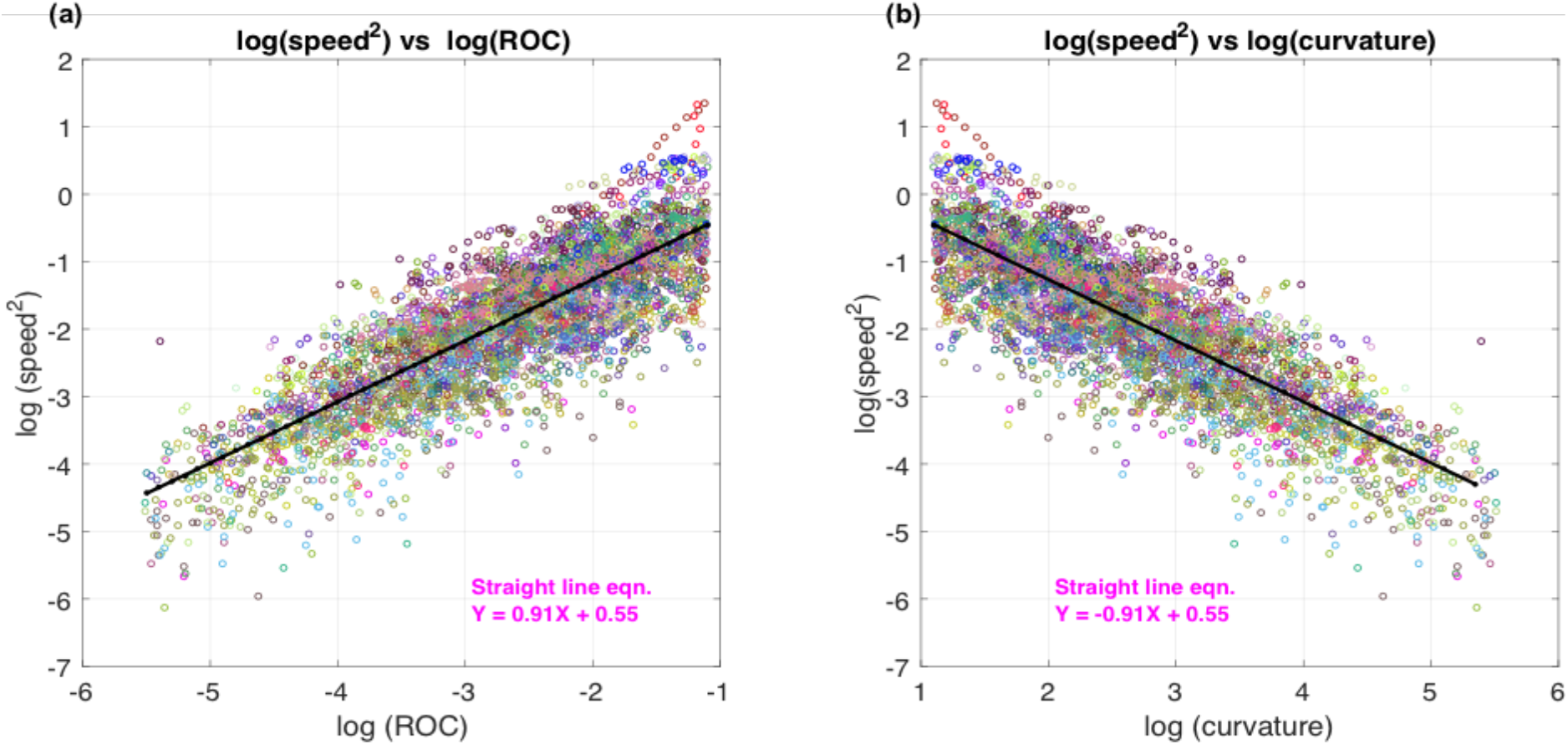
*Scatterplot of log(speed^2^) vs log(ROC) (b): Scatterplot of log(speed^2^) with log(curvature) for 64 bee trajectories*.

Fig. 9b shows the relationship between log(speed^2^) and log (curvature). This relationship is again approximately linear, with a slope of −0.9, which is close to the value of-1.0 predicted by the hypothesis.

The Y-axis intercept of the regression lines shown in Fig. 9a and 9b is 0.55, from which the average centripetal acceleration can be calculated to be *e*^0.55^ = 1.73 m/s^2^. This is not very different from the value of 1.99 m/s^2^ estimated from the slope of the regression of the data in Fig. 8a, the difference arising because of the scatterplot in Fig. 8a is transformed nonlinearly to obtain the scatterplots of Figs. 9a and 9b.

Computing the slope of the regression line of a scatterplot of the relationship between log (speed) and log (radius of curvature) yields a value of 0.45 (plot not shown). This value is very close to the expected value of 0.5, additionally supporting our hypothesis of constant centripetal acceleration.

Thus, broadly speaking, the data of figs. 8 and 9 support the hypothesis that bees maintain a more or less constant centripetal acceleration during their turns, irrespective of the instantaneous speed or curvature at each point along the turn.

Table 1 compares the coefficients of variation (CV) of the variables that characterise the trajectories. We observe that, although the CV of the curvature is relatively high, signifying relatively large variations in curvature magnitude, the CV of the centripetal acceleration magnitude is relatively low. This is because the bees are tailoring the flight speed to the curvature - reducing speed during tight turns, and increasing it during shallow turns – thus reducing variations in the CA. Although the variations in flight speed are relatively low (CV=0.29), the variations in speed^2^ are comparatively high (CV= 0.57). This variation works to counteract variations in the centripetal acceleration that could potentially arise from variations in the curvature if the speed were held constant. The reciprocal interaction between these two variables is evident from Eqn. (7).

**Table 1:** Mean coefficients of variation (CV) of curvature, speed, speed^2^, and centripetal acceleration magnitude

| Parameter | Curvature | Speed | Speed<sup>2</sup> | Centripetal acceleration magnitude |
| --- | --- | --- | --- | --- |
| CV | 0.89 | 0.29 | 0.57 | 0.42 |

### Comparison of characteristics of left and right turns

We were interested to examine whether the bees showed any preferences for turning direction. If the bee’s rotation about its dorsoventral axis (Zn) is in the clockwise direction, then the bee turns to the right, and vice versa. In order to determine the turning direction, we computed the 3D rotation vector, which is given by the cross product between the unit velocity vector and the unit centripetal acceleration vector (see Section III of the SM for explanation). The bee’s turning direction is then obtained by taking the dot product of the 3D rotation vector with the unit vector representing the dorsoventral axis of the bee. If the dot product is positive, the bee is turning right; and vice versa.

This procedure was used to classify the turning direction, and then to compare the curvatures and centripetal accelerations during left turns with those during right turns. The histogram in Fig. 10 compares the distributions of the curvatures of right turns with those of left turns. Positive curvatures represent right turns, and negative curvatures left turns. The histogram is nearly symmetrical. The mean curvature magnitudes during left (-19.3 m^−1^) and right (17.2 m^−1^) turns are more or less equal and not significantly different (*p* = 0.230; two tailed t-test). The overall mean curvature for all turns (−0.97 m^−1^) is very close to zero and is not significantly different from zero (*p* = 0.277; two tailed t-test). This indicates that turns in either direction are (a) equally likely, and (b) display the same distribution of curvature magnitudes. Thus, the bees flying in our experimental situation do not display any noticeable left-right biases in their turning behaviour.

**Figure 10:**
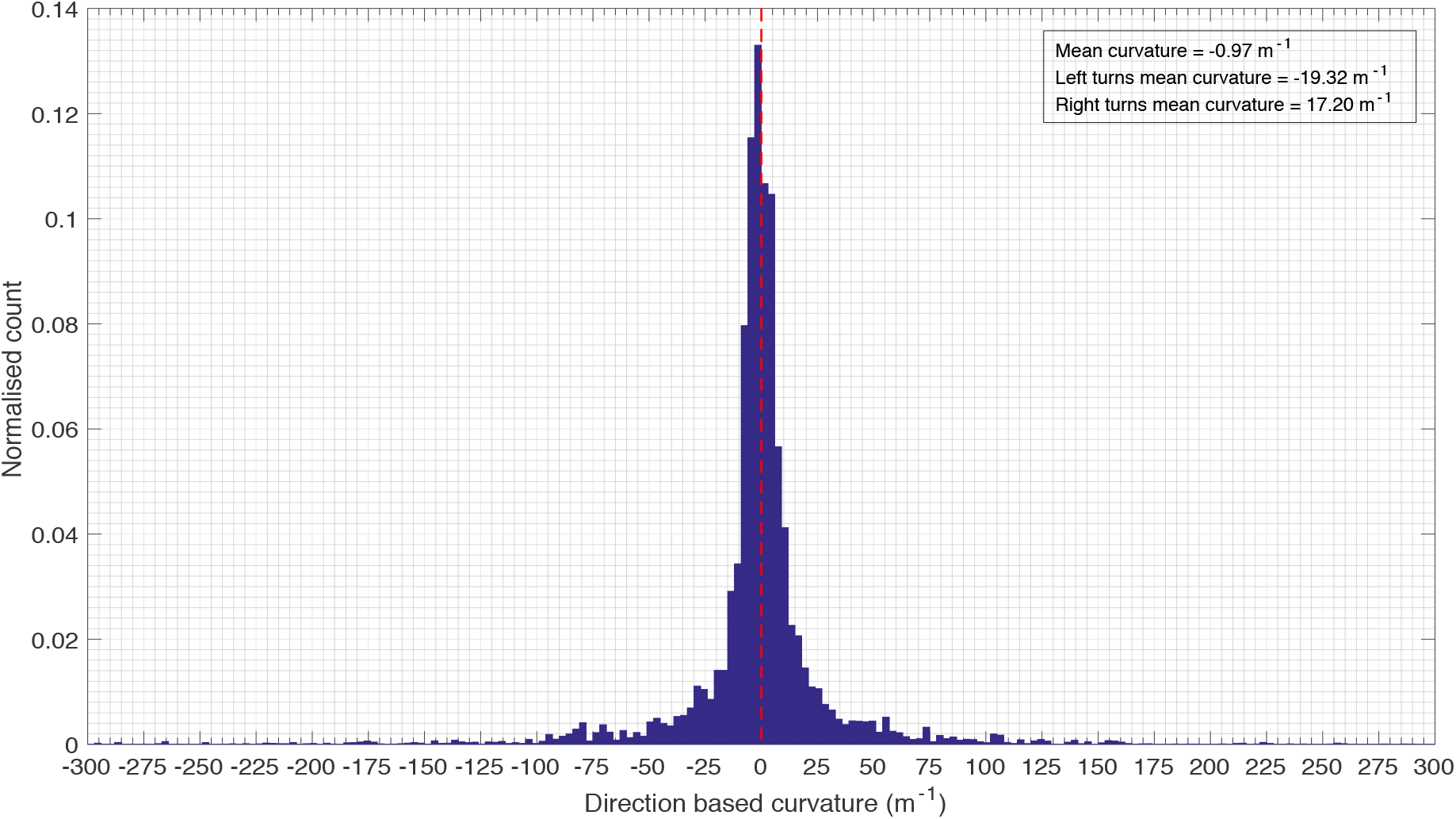
*Normalised histogram of direction based curvature for 64 bees*.

We also looked for possible biases in the centripetal accelerations associated with left versus right turns. Fig. 11 shows a histogram of the distribution of centripetal acceleration for all bees. The peak value of the centripetal acceleration is slightly higher for left turns than for right turns. Apart from this, the histograms for the left and right turns are nearly symmetrical - the mean centrifugal acceleration for right turns (+2.5 m/s^2^) is not significantly different from that for the left turns (−2.7 m/s^2^; *p* = 0.460, two tailed t-test). The overall mean centrifugal acceleration for all turns (−0.28 m/s^2^) is very close to zero and is not significantly different from zero (*p* = 0.683; two tailed t-test). Thus, as with the curvature magnitudes, there is no major overall bias in the distribution of the centripetal accelerations.

**Figure 11:**
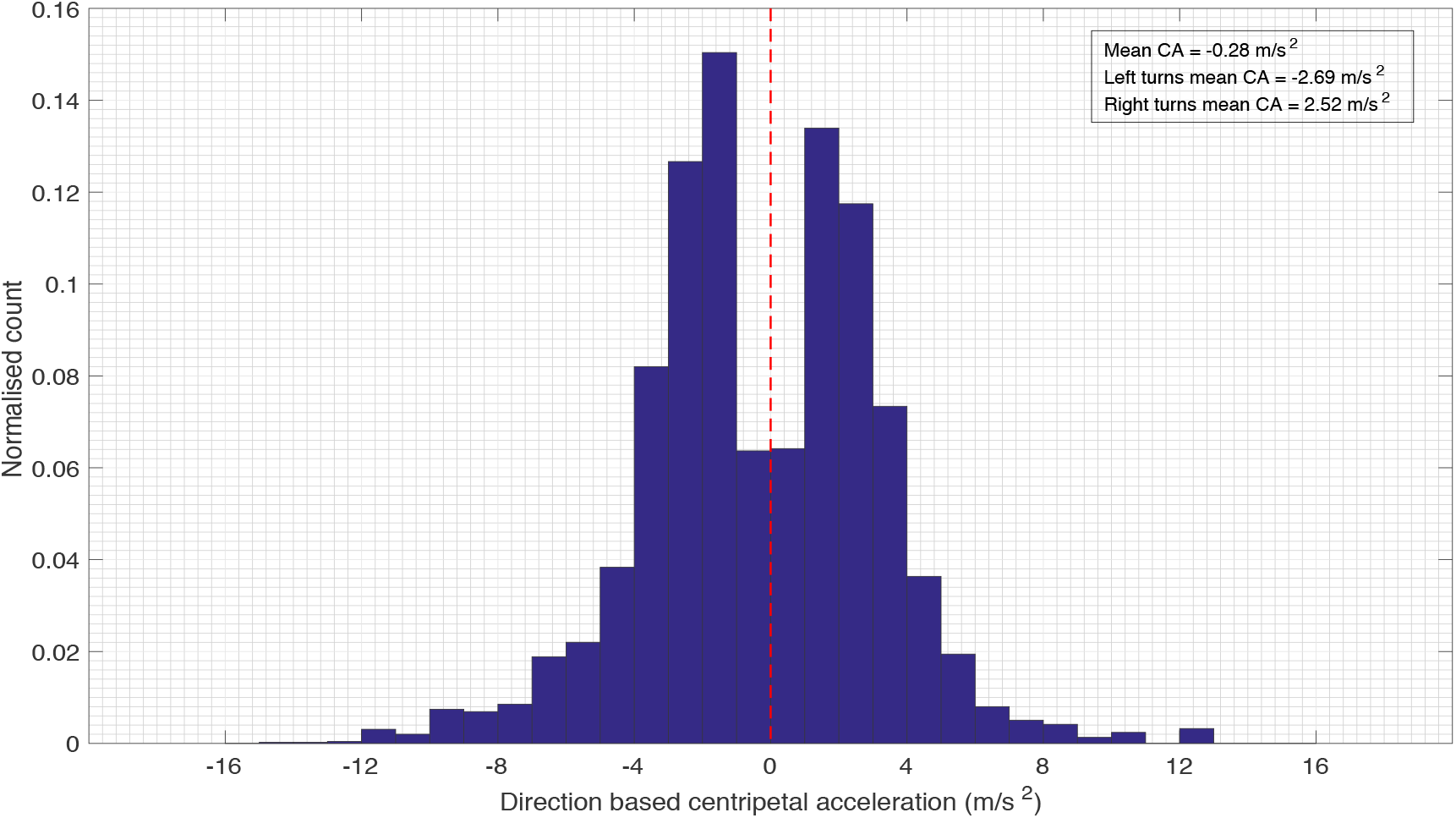
*Normalised histogram of direction based centripetal acceleration for 64 bees*.

### Body deviation angle analysis of turning bees

From the analysis presented above, we have hypothesized that bees keep their centripetal acceleration almost constant during turns. This strategy might help them perform coordinated turns, without deviating from the intended flight trajectory. Accordingly, we were interested to look for evidence of sideslip. This was done by examining the body deviation angle (BD angle) during turns. We define the BD angle as the angle, measured in the horizontal plane, between the instantaneous flight direction vector and the instantaneous bee’s body orientation vector. This angle is zero when the body axis is aligned with the flight direction. Its polarity is defined to be negative when the body axis points into the turn, and positive when the body axis points away from the turn. We commenced the analysis by calculating the BD angles and plotting their histograms during left turns, right turns and straight flight. “Straight flights” were defined to be sections of the trajectory in which the curvature magnitude was lower than a threshold of 5.0, and turning flights were sections in which the curvature magnitude exceeded 5.0, with the polarity of the curvature defining the direction of the turn. The results are shown in Figs. 12–14, where each histogram has been fitted to a Gaussian distribution.

**Figure 12:**
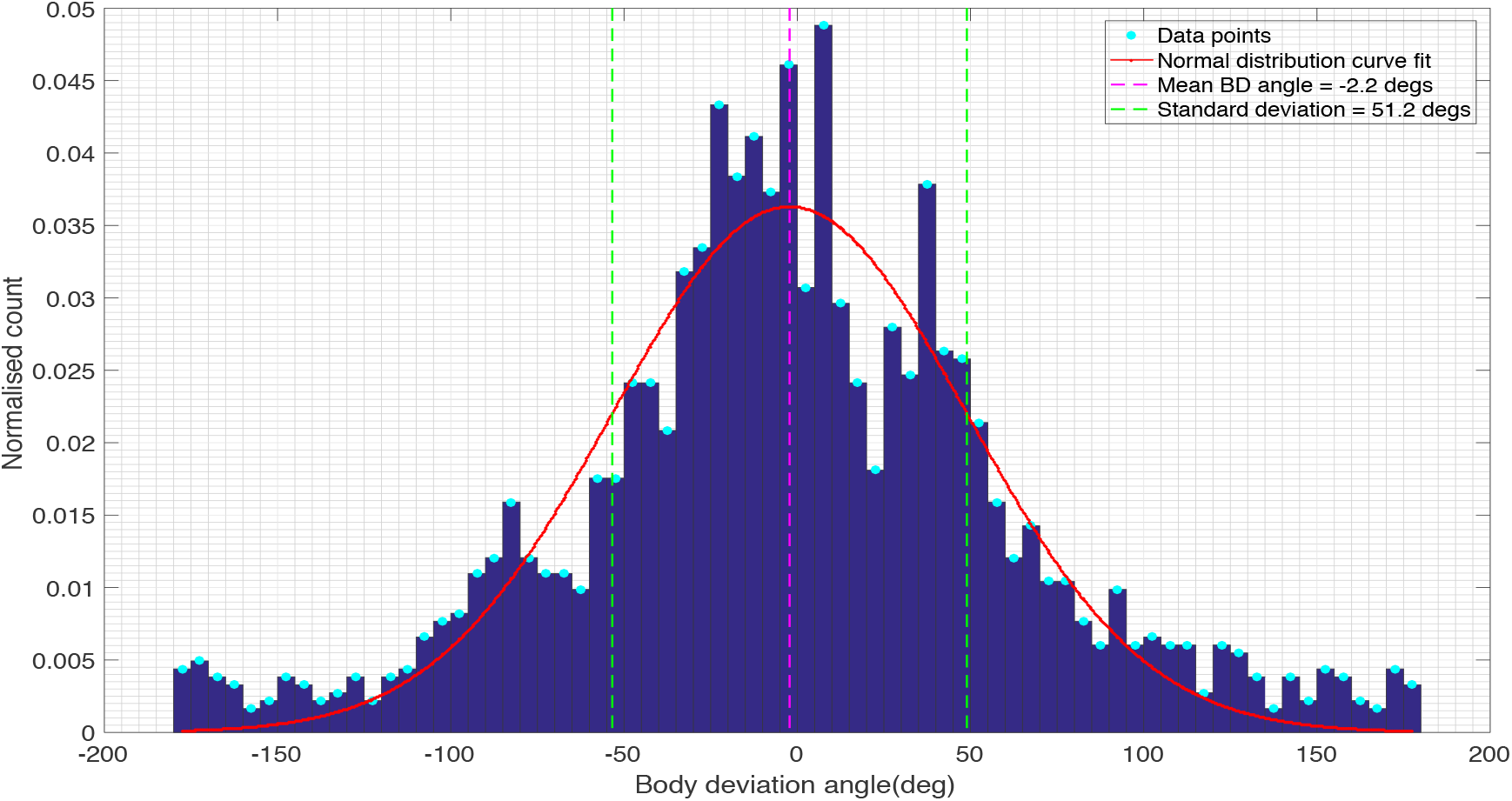
*Histogram of BD angles during left turns, fitted to a Gaussian distribution*

**Figure 13:**
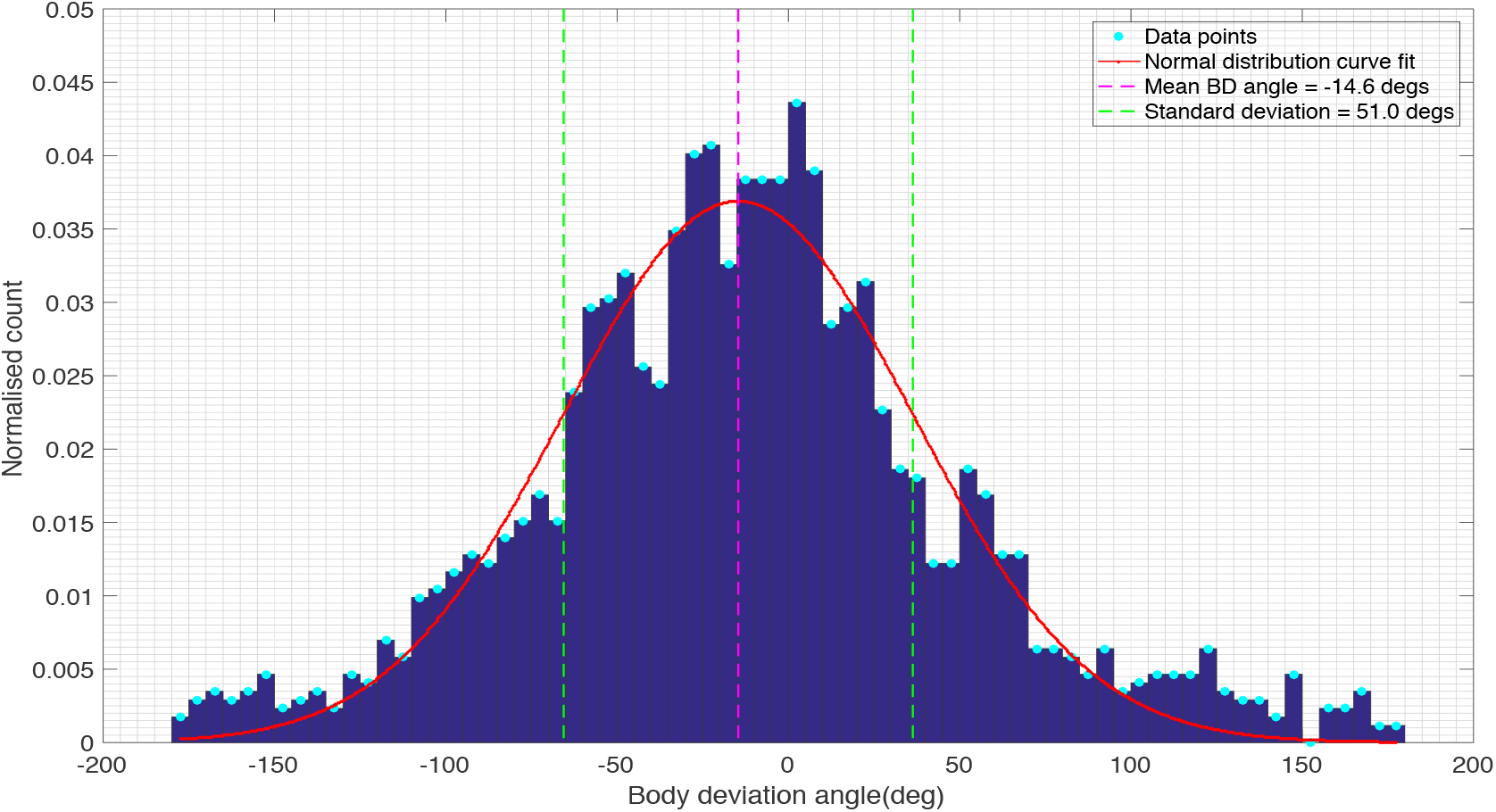
*Histogram of BD angles during right turns, fitted to a Gaussian distribution*.

**Figure 14:**
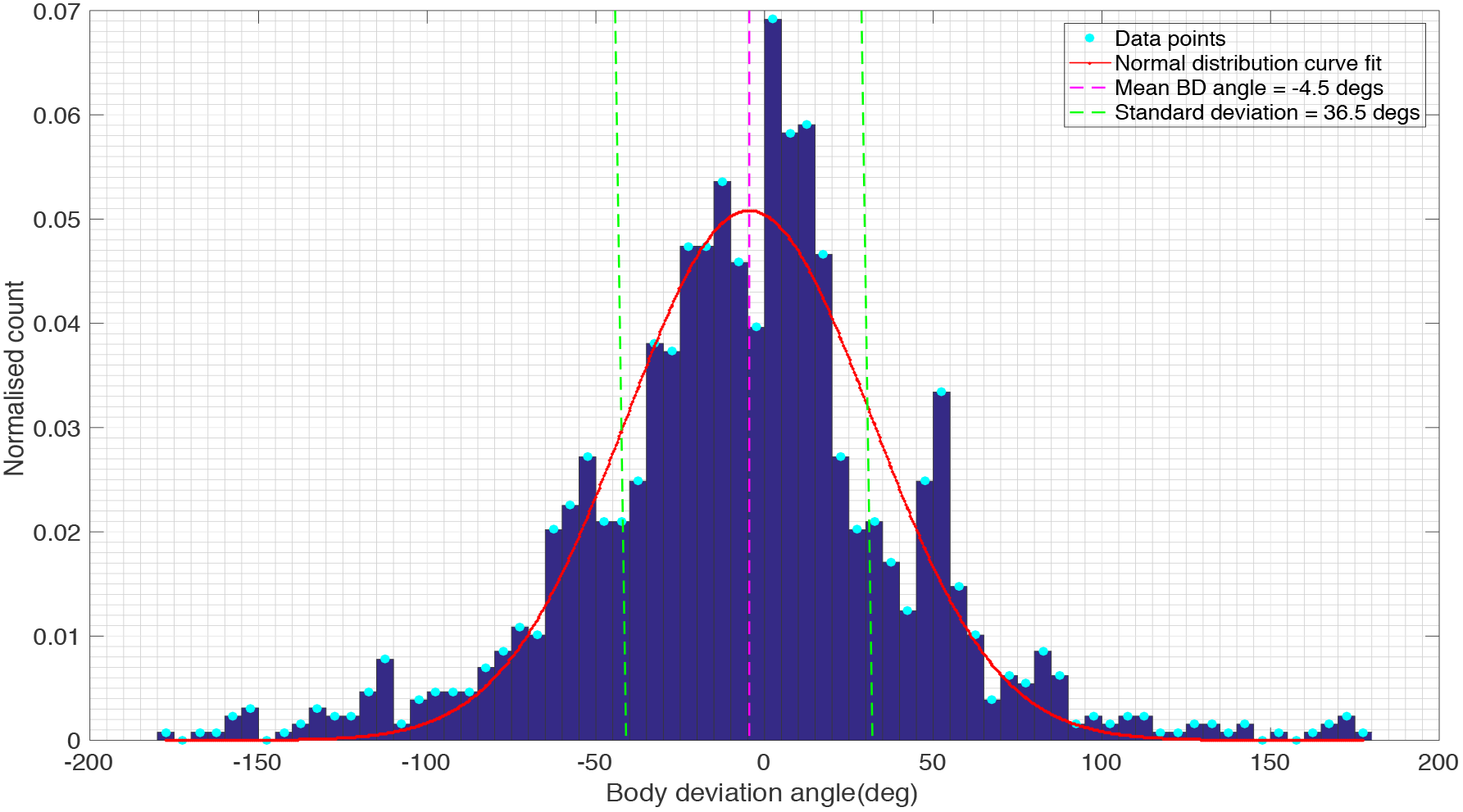
*Histogram of BD angles during straight flights, fitted to a Gaussian distribution*.

Table 2 gives values of the mean and standard deviation of the body deviation angle after correction for estimated errors in the measurement of the direction of body orientation and flight direction from the video images, as described in Section IV of the SM. The results reveal that the BD histograms for left turns, right turns and straight flight display a mean value close to zero, but a broad standard deviation of about 50 deg. This implies that, although the body orientation can occasionally deviate substantially from the direction of flight, the deviations are more or less symmetrical, with roughly half of the deviations pointing into the turn and the other half pointing outward; and this is true for all three conditions - left turns, right turns and even in straight flight. This suggests that the observed BDs are not a reflection of uncontrolled turns that involve sideslips; rather, they are a natural characteristic of the loitering bees, in which the body does not point consistently in the flight direction. Sideslips, if present, would be reflected in the left and right-turn histograms by an increased frequency of negative BD angles (body pointing into the turn) - which is not the case. Instances where the magnitude of the BD angle exceeds 90 deg represent situations where the bee is moving temporarily backwards. In this condition, the BD angles are again positive or negative equally frequently.

**Table 2:** Mean, raw standard deviation and corrected standard deviation of BD angle

| Flight category | Mean BD angle (deg) | Raw standard deviation of BD angle (deg) | Corrected standard deviation of BD angle (deg) |
| --- | --- | --- | --- |
| Left turns | -2.2 | 51.2 | 48.2 |
| Right turns | -14.6 | 51.0 | 48.0 |
| Straight flights | -4.5 | 36.5 | 32.3 |

To further explore the existence of sideslips, we computed the mean value of the magnitude of the centripetal acceleration in each bin of the body deviation angle histograms of Figs. 12–14. The results, shown in Figs. 15–17, indicate that the magnitude of the centripetal acceleration is more or less constant, independent of body deviation angle. This is true for right turns, left turns, and straight flights. BD angles greater than 90 deg are not included in the histogram (Figs. 15–17), since those instances are not turning flights, but rather situations in which the bee is moving temporarily backwards.

**Figure 15:**
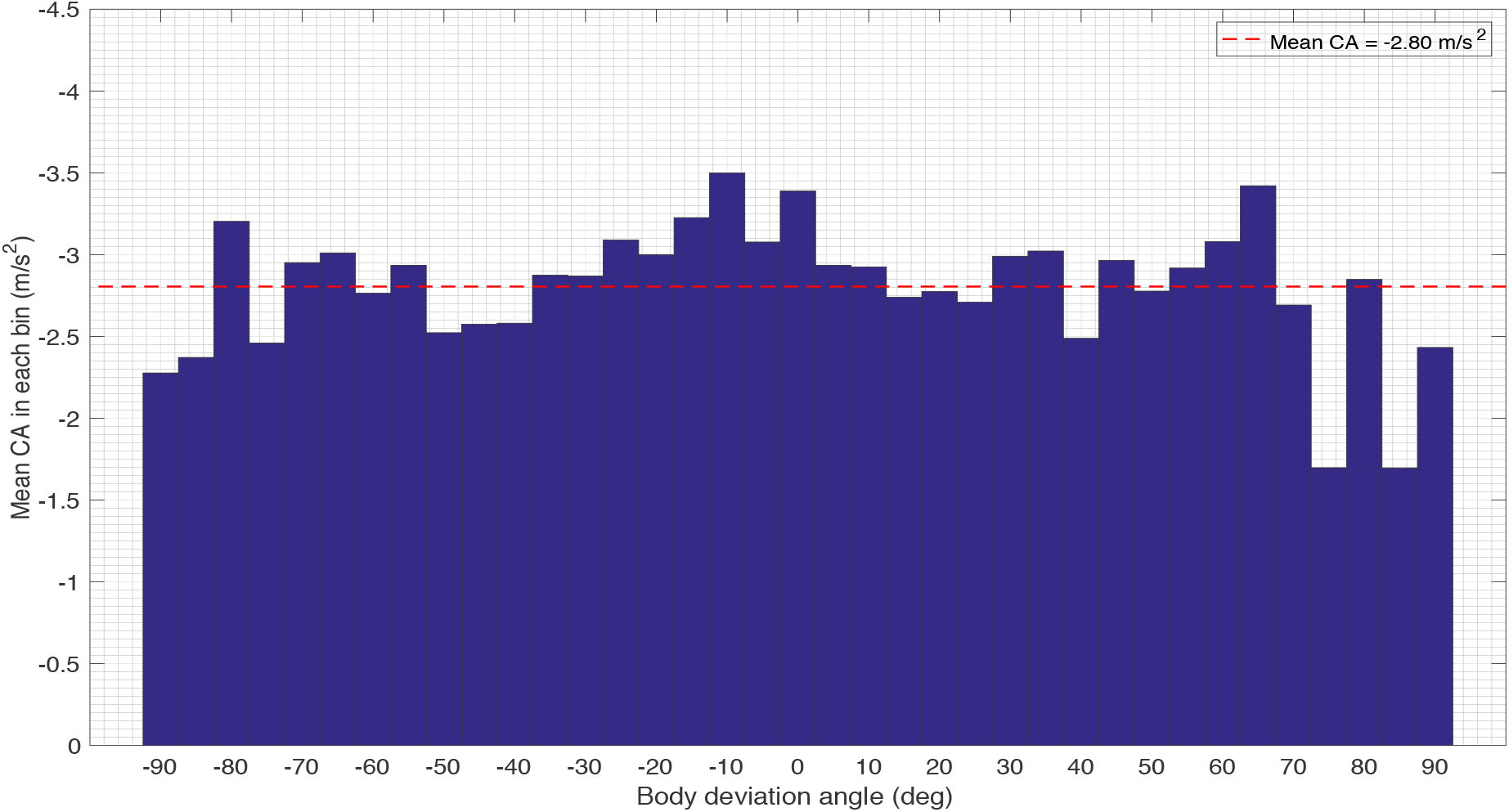
*Variation of mean centripetal acceleration magnitude with BD angle during left turns. The dashed red line represents the overall mean*.

**Figure 16:**
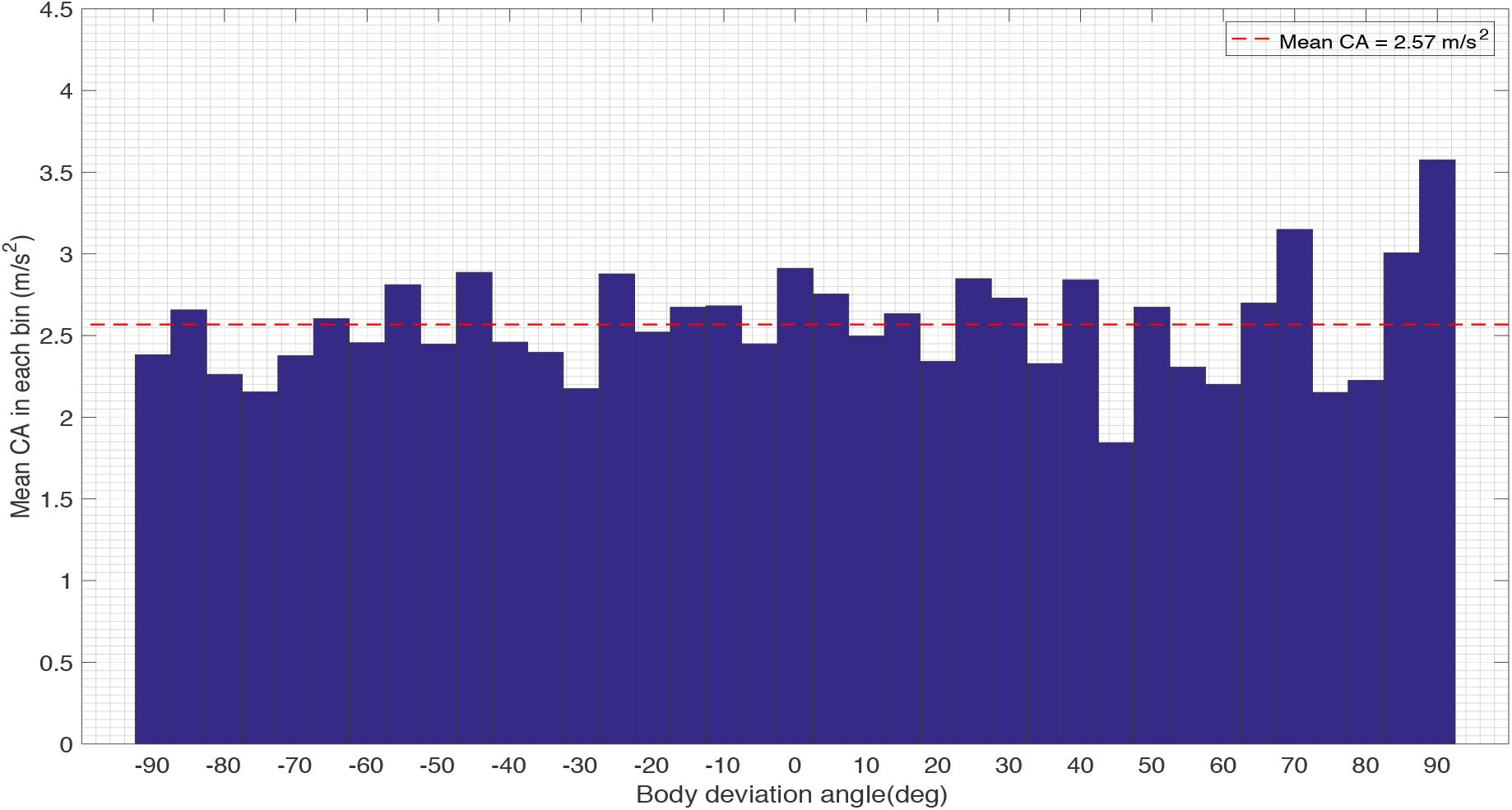
*Variation of mean centripetal acceleration magnitude with BD angle during right turns. The dashed red line represents the overall mean*.

**Figure 17:**
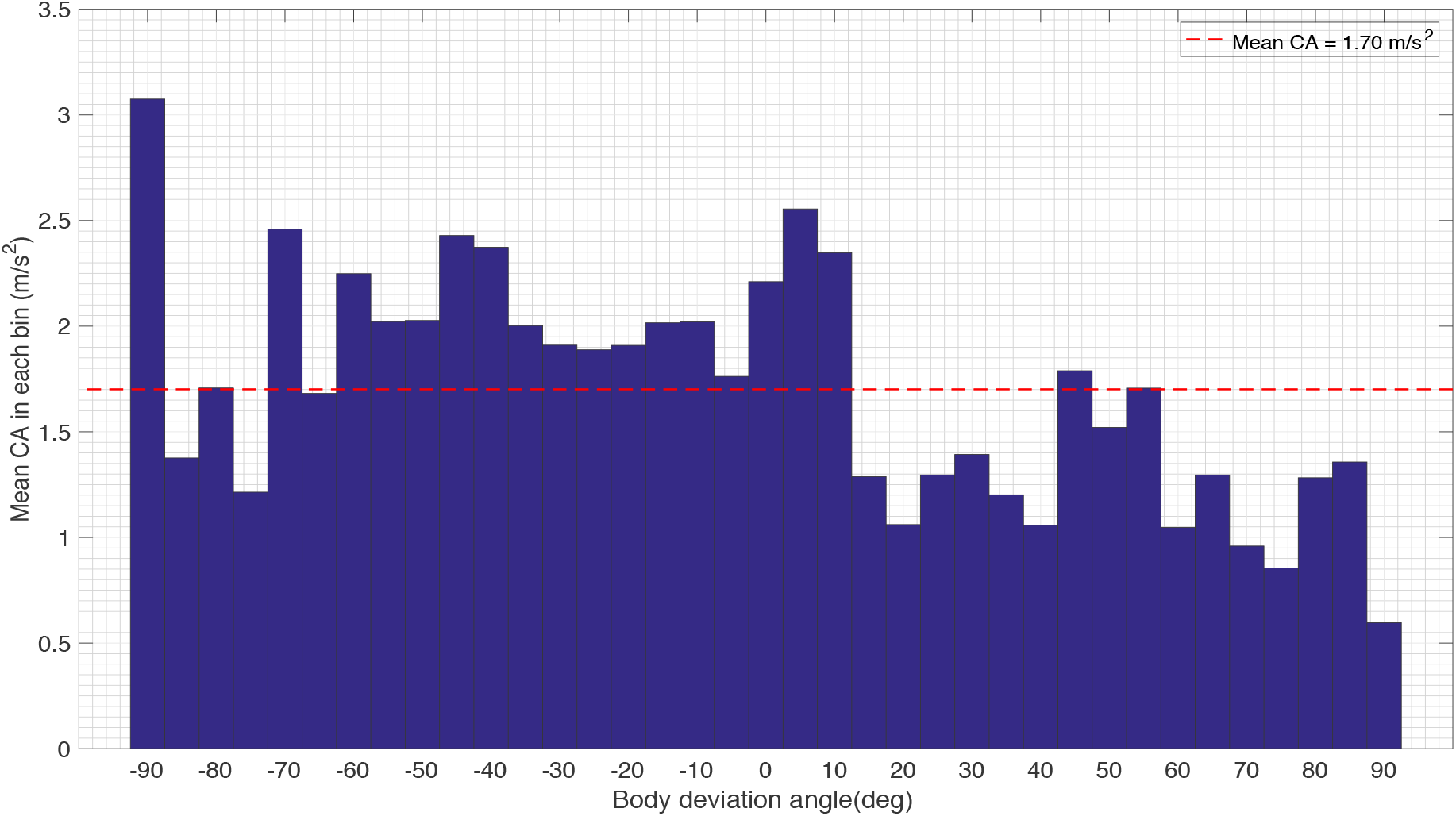
*Variation of mean centripetal acceleration magnitude with BD angle during straight flights. The dashed red line represents the overall mean*.

The mean value of the CA magnitude, computed from the histograms of Figs. 15–17, are −2.80 m/s^2^ for left turns, 2.57 m/s^2^ for right turns, and 1.70 m/s^2^ for close-to-straight flights.

Secondly, the observation that the CA magnitudes are similar even at large negative and positive values of BD angle (i.e. irrespective of whether the body is pointing sharply into or away from the turn), makes it very unlikely that the large negative values of BD angles (when the body is pointing sharply into the turn) are associated with sideslips or skids. In summary, the data in Figs. 12–17 and Table 2 suggest that the bees flying in the cloud are never overcome by the centrifugal forces that are encountered while executing these turns, which would result in sideslips.

## DISCUSSION

We have investigated the turning flight characteristics of loitering honeybees in a semi-outdoor environment comprising a number of bees flying in close proximity to each other, trying to enter a blocked hive. We commenced our analysis by studying how the kinematics of bees vary in a cloud. In general, the speed of the bee varies continuously through its flight path. We observed that the flight speed tends to decrease whilst entering a turn, and increase whilst exiting it. This was indicated by the direction of the tangential acceleration vector, which points against the flight direction while entering the turn, and in the flight direction when leaving it.

This general pattern of speed variation has been documented in a number of other aerial and terrestrial animal species, for example fruit flies (Mronz and Lehmann, 2008), bats (Aldridge, 1987), horses (Tan and Wilson, 2011) and northern quolls (Wynn et al., 2015). However, none of these studies have quantitatively examined the relationship between speed and turning radius. The present study does this and finds that, during the course of a turn, flight speed varies with curvature in such a way that the centrifugal force is maintained at a more or less constant value, irrespective of the moment-to-moment variations in speed and curvature.

Our results provide an estimate of this centrifugal force. The histogram of Fig. 11 indicates that the mean centripetal acceleration is −2.69 m/s^2^ during left turns, and 2.52 m/s^2^ during right turns. This is in good agreement with the data shown in Figs. 15 and 16, which indicate mean centripetal accelerations of −2.80 m/s^2^ for left turns, and 2.57 m/s^2^ for right turns. The values from the two analyses are not exactly the same for the two analyses because body deviation angles greater than 90 deg were excluded from the histograms of Figs. 15–17. Overall, our results and analysis reveal that the magnitude of the centrifugal force does not exceed 2.8 m/s^2^, or 0.29g, where *g* is the acceleration due to gravity. Thus, we infer that the centrifugal force is not greater than about 30% of the bee’s weight, which we propose is low enough to permit coordinated turns without incurring unwanted sideslips. Orchestrating turns in this way would ensure that the insect is never dominated by the centrifugal force during the turn, and always maintains the intended (curved) trajectory.

To probe our hypothesis further, we examined whether bees undergo sideslips during turns. If a bee is unable to resist the centrifugal force that it experiences during a turn, we would expect its body to point into the turn - analogous to a car that skids out of control while making a sharp turn. Our findings (Figs. 12–14) indicate that there is no systematic bias in the body deviation angle that is correlated with the direction of the turn - in other words, there is no evidence that the body points preferentially into the turn. Moreover, the finding that the width of the body deviation histogram is approximately the same for left turns, right turns and nearly straight flights (see Table 1), suggests that the variations in the body deviation angle are a normal feature of honeybee flight in these experimental conditions, and do not reflect sideslips. Additional evidence for the lack of sideslips comes from the plots of centripetal acceleration versus body deviation angle (Figs. 15–17), which reveal that the centripetal acceleration is roughly constant – it does not vary with the body deviation angle. If sideslips were to occur, one would expect large body deviations into the turn (negative body deviation angles) to be associated with larger centrifugal accelerations. This is clearly not the case – there is no correlation between the body deviation angle and the centripetal acceleration (or, equivalently, centrifugal force) – which, again, suggests that the observed variation in the body deviation angles is not due to the presence of uncontrolled turns.

Finally, our study also indicates that, under our experimental conditions, left and right turns display similar characteristics, when the data are pooled across the bees that were investigated. Thus, it appears that, as a whole, the group of bees flying in our bee cloud does not exhibit a preferred turning direction (left or right) - although we cannot rule out the possibility that individual bees have turning biases, which would be a topic for future investigation. On the other hand, army ants, fish (Vicsek and Zafeiris, 2012) and bats (Ren et al., 2016) rotate in a particular direction displaying a collective turning behaviour that could promote collision avoidance. In summary, our study documents a turning strategy that is used by honeybees to execute controlled, side-slip free turns while they are in a loitering mode of flight in a bee cloud. It would be interesting to examine whether this strategy also applies to flight in other conditions.

## ACKNOWLEDGEMENTS

We thank Prof. Thomas Stace from the Department of Physics, University of Queensland, for providing valuable suggestions for analysis of the data. Our special thanks to Mr. Peter Anderson, for the care of the bee colony used in this experiment. This study was supported by a UQ International student scholarship and a Boeing top-up scholarship awarded to MYM, and Australian Discovery Research Grant DP 140100914 to MVS.

